# *C. elegans* genetic background modifies the core transcriptional response in an α-synuclein model of Parkinson’s disease

**DOI:** 10.1101/348623

**Authors:** Yiru A. Wang, Basten L. Snoek, Mark G. Sterken, Joost A.G. Riksen, Jana J. Stastna, Jan E. Kammenga, Simon C. Harvey

## Abstract

Accumulation of protein aggregates is a major cause of Parkinson’s disease (PD), a progressive neurodegenerative condition that is one of the most common causes of dementia. Transgenic *Caenorhabditis elegans* worms expressing the human synaptic protein α-synuclein show inclusions of aggregated protein and replicate the defining pathological hallmarks of PD. It is however not known how PD progression and pathology differs among individual genetic backgrounds. Here, we compared gene expression patterns, and investigated the phenotypic consequences of transgenic α-synuclein expression in five different *C. elegans* genetic backgrounds. Transcriptome analysis indicates that the effects of α-synuclein expression on pathways associated with nutrient storage, lipid transportation and ion exchange depend on the genetic background. The gene expression changes we observe suggest that a range of phenotypes will be affected by α-synuclein expression. We experimentally confirm this, showing that the transgenic lines generally show delayed development, reduced lifespan, and an increased rate of matricidal hatching. These phenotypic effects coincide with the core changes in gene expression, linking developmental arrest, mobility, metabolic and cellular repair mechanisms to α-synuclein expression. Together, our results show both genotype-specific effects and core alterations in global gene expression and in phenotype in response to α-synuclein. We conclude that the PD effects are substantially modified by the genetic background, illustrating that genetic background mechanisms should be elucidated to understand individual variation in PD.

## Introduction

Dementia is a growing global problem, and as life expectancy increases so does the number of people affected. The most common causes of dementia are associated with neurodegeneration, such as that resulting from Alzheimer’s disease (AD) and Parkinson’s disease (PD) (Kivipelto *et al.* 2005; Aarsland & Kurz 2018; Aybek *et al.* 2009). Despite their distinct causes, these neurodegenerative dementias share neuroanatomical and biochemical similarities, both result from protein misfolding, and both show overlapping cognitive and behavioural symptoms.

A major difficulty in determining the mechanisms that produce neurodegenerative dementia is that the underlying cellular-level pathology is highly variable between individuals (reviewed in Seelaar *et al.* 2008; Gratwicke *et al.* 2015). This results from background modifiers affecting cellular pathways that influence disease onset and progression. The consequence of this is substantial variability in age of onset and disease progression. This complicates attempts to develop both diagnostic tests and treatment options at the individual level.

Model organisms, such as the nematode *Caenorhabditis elegans*, are of great value for studying the genetics of complex human diseases (Teschendorf & Link 2009; Calahorro & Ruiz-Rubio 2011; Rodriguez *et al.* 2013; Sin *et al.* 2014; Wang *et al.* 2017). This is due to their experimental tractability and to the broad and general conservation of genetic pathways between species. Importantly, *C. elegans* allows for sophisticated genome-wide genetic screens, with most such studies either seeking to identify loci that represent candidate diagnostic and therapeutic targets, or to understand the underlying pathological processes. For example, analysis of transgenic *C. elegans* expressing the human Aβ peptide - the main component of the amyloid plaques found in AD - linked toxicity to insulin/insulin-like growth factor (IGF), dietary restriction and the heat shock response (Fonte *et al.* 2008; Steinkraus *et al.* 2008; Teschendorf & Link 2009; Calahorro & Ruiz-Rubio 2011; McColl *et al.* 2012; Sin *et al.* 2014). Similarly for PD, analysis of worms expressing α-synuclein identified associations with ageing and insulin-like signalling (van Ham *et al.* 2008).

Most *C. elegans* studies have however been undertaken on only the canonical Bristol N2 genetic background, and therefore do not provide insight into how genetic variation between individuals might affect PD-associated traits. This is a major issue as it is clear, both specifically in *C. elegans (e.g.* Duveau & Félix 2012; Elvin *et al.* 2011; Schmid *et al.* 2015; Sterken *et al.* 2017) and more generally in other systems (Wade & Daly 2005; Justice & Dhillon 2016; Coe *et al.* 2009), that the phenotype of any given mutation, transgene or allele can vary depending on the genetic background (Kammenga 2017). For *C. elegans*, natural genetic variation is known to result in extensive phenotypic variation *(e.g.* Reynolds & Phillips 2013; Martin *et al.* 2017 and see Kammenga *et al.* 2008 and Gaertner & Phillips 2010 for reviews for older studies) and differentially affects both the proteome (Kamkina *et al.* 2016; Singh *et al.* 2016) and transcriptome (Li *et al.* 2006; Rockman *et al.* 2010; Viñuela *et al.* 2010; Volkers *et al.* 2013; Kamkina *et al.* 2016; Snoek *et al.* 2017; Sterken *et al.* 2017).

Exemplifying this pattern of a reliance on a single genetic background, only one study has, to date, looked at protein misfolding disease in multiple wild isolate genetic backgrounds of *C. elegans* (Gidalevitz *et al.* 2013). Crucially, this showed that natural genetic variation can uncouple different phenotypic effects of polyglutamine (polyQ40) expression (Gidalevitz *et al.* 2013). This strongly suggests that similar important variation between genetic backgrounds will be found for other protein misfolding diseases. Given that for protein misfolding diseases the nature of the modifying alleles segregating within human populations remain largely elusive (Kearney 2011), the experimental tractability of *C. elegans* makes the species an excellent system in which to address this issue.

In humans, α-synuclein plays a central role in the aetiology and progression of PD (Schulz-Schaeffer 2010). An increase in the α-synuclein gene dosage causes early-onset forms of familial PD in humans (Singleton *et al.* 2003). In *C. elegans*, expression of an α-synuclein and yellow fluorescent protein (YFP) fusion in the body wall muscle results in an age-dependent accumulation of inclusions (van Ham *et al.* 2008). These inclusions of α-synuclein form aggregates in aging worms that are similar to the pathological inclusions seen in humans with PD (van Ham *et al.* 2008). To investigate the effect of genetic background on PD, we have created introgression lines (ILs) containing this α-synuclein and YFP transgene in the background of four wild isolates of *C. elegans.* Our analyses of these new ILs, and of α-synuclein in an N2 genetic background, identify both general and genotype-specific changes in gene expression. These changes predict a range of phenotypic effects that we then experimentally confirm. Importantly, given the reliance on N2 in *C. elegans* research, we show both that some effects are N2-specific and that for other phenotypes the analysis of other genetic backgrounds uncovers substantial variation not seen in N2.

## Results

### Introgression line construction and validation

We introgressed the *pkIs2386* transgene [unc-54p:: α-synuclein::YFP + unc-119(+)] from NL5901 (van Ham *et al.* 2008), which has an N2 genetic background, into the genetically distant wild isolates JU1511, JU1926, JU1931, and JU1941 (see Volkers *et al.* 2013 for information on genetic distance between these lines). After back-crossing and selfing, four new ILs were obtained, SCH1511, SCH1926, SCH1931, and SCH1941, with for example, SCH1511 containing the transgene in a JU1511 background. In combination with N2 and NL5901 as controls, we were therefore able to investigate the phenotypic and genomic effects of α-synuclein in five genetic backgrounds (N2, JU1511, JU1926, JU1931, and JU1941).

We sought to identify the site of the introgression and to determine how much of the N2 genome surrounding the transgene had also been introgressed into the wild isolates. PCR-based genotyping located the transgene in chromosome IV, indicating that the homozygous introgressions in the new ILs spanned between 4.2 and 13.2Mb of chromosome IV (Table S1). By using the set of genetic markers from Volkers *et al.* (2013), we identified significantly differential expressed genes on chromosome IV of the PD lines, and also identified an additional introgression on chromosome V in SCH1931 (Figure S1). PCR-based genotyping together with the transcriptomic analysis indicated a consistent genomic location for the PD transgene in each of the genetic backgrounds, but did not allow detection of the precise introgression boundaries.

### Transcriptional response to PD in different genetic backgrounds

Initial analysis of genome-wide transcriptional changes in α-synuclein introgression worms suggested differences in developmental rate between ILs. Variation between ILs in developmental rate was also observed during IL construction. We therefore estimated the age of the samples by their gene expression profile (as in Snoek *et al.* 2014a and van der Bent *et al.* 2014) and used principal component analysis (PCA) on genome-wide expression levels to investigate differences between isolates and the effects of α-synuclein. This analysis revealed separation between the wild isolates and their corresponding transgenic ILs (Figure 1A). Although genotypes were more scattered on these first two PCA axes, this indicated that developmental delay and reduced lifespan were associated with the introgressed PD transgene (Figure 1A). A large part of the global gene expression differences are therefore explained by slower development in the PD lines. Hence, many of the genes induced in response to α-synuclein in different backgrounds were likely to be associated with their differential age. Using a threshold of -log10(p) > 3.4 (FDR < 0.05) we found 3521 genes affected by Age, 1509 affected by Genotype, 646 by PD and 428 by an interaction between Genotype and PD (Figure 1B, Table S2). Most differentially expressed genes are affected by age alone (2893) or only by genotype (815). Genes affected by PD or the interaction between PD and age were most often also affected by genotype. The overlap among these four factors yielded 79 genes highly specific for α-synuclein relative to both

**Figure 1:**
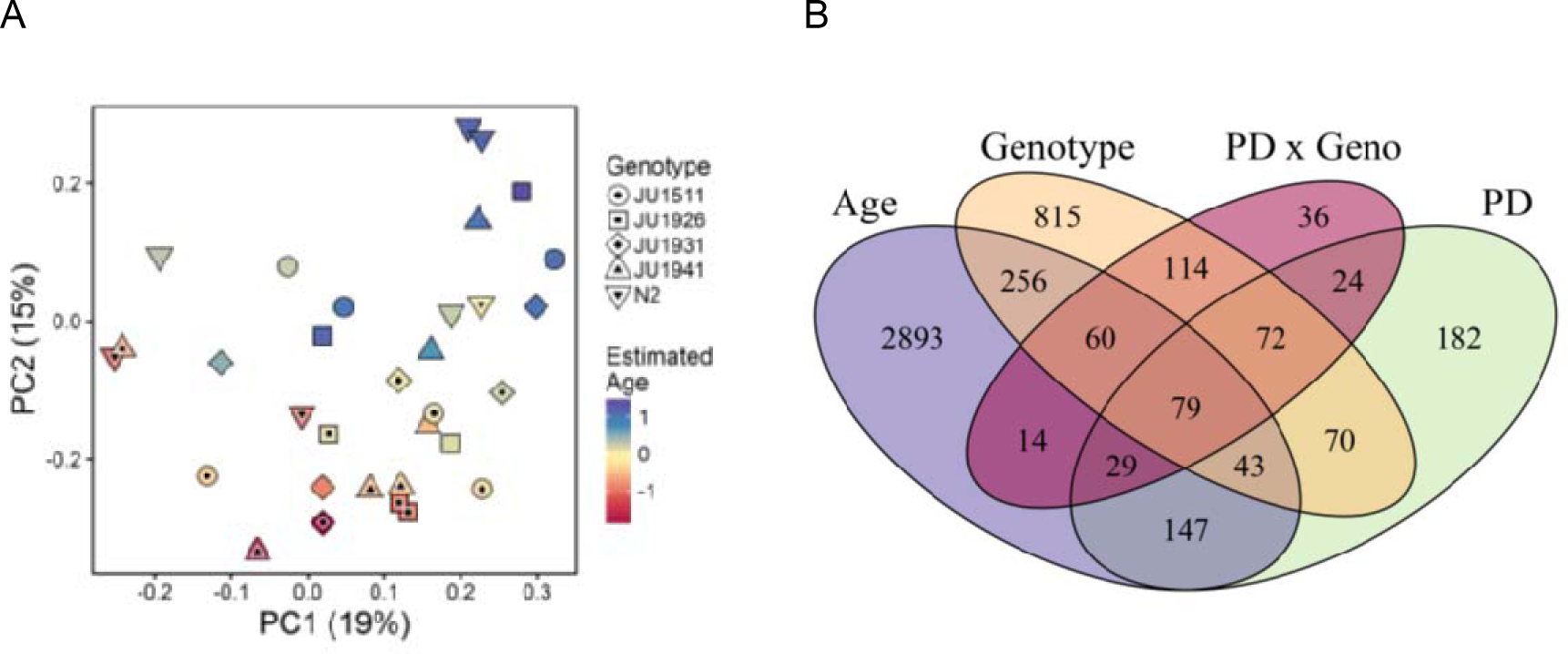
**A)** PCA on gene expression differences. Estimated age difference is shown by the colour gradient, PD background shown by black point. B) Venn diagram of differentially expressed genes that are specific to PD, Genotype, Age and PD x Geno including both induced and repressed genes.

Age and Genotype (Table S2). This showed that both genotype-specific and universal (genotype independent) changes in gene expression were induced by α-synuclein expression, *i.e.* we identified a core set of genes of which the expression was altered in all genetic backgrounds and others that were genotype-specific. Strikingly, many genes were not expressed and/or regulated in the same way in wild isolate genetic backgrounds compared with N2. These data therefore indicate that the transcriptome effects of the α-synuclein introgression depend on the genetic background.

Gene ontology analysis (GO) of differentially expressed genes revealed the molecular, cellular, and biological processes affected by development, PD and genotype (Table 1, with full results in Table S2). Genes that change expression in response to genotype were enriched for genes involved in the innate immune response and oxidation-reduction process (Table S2) as was previously found by Volkers *et al.* (2013). Widespread changes in genes related to muscle function were observed (Table 1), an expected response given the changes in cellular environment induced by expression of **α**-synuclein in the body wall muscle. This analysis also identified changes in genes involved in pharyngeal pumping (Table 1).

**Table 1:** The top-20 enriched GO-terms with 5 or more genes with a PD effect.

| GO_ID | Description | All Genes | PD Genes | Type | *p*-value | adjusted *p* |
| --- | --- | --- | --- | --- | --- | --- |
| GO:0005576 | extracellular region | 329 | 54 | CC | 3.99E-06 | 8.29E-04 |
| GO:0030246 | carbohydrate binding | 288 | 48 | MF | 8.12E-06 | 1.13E-03 |
| GO:0005861 | troponin complex | 8 | 5 | CC | 1.21E-05 | 1.26E-03 |
| GO:0055120 | striated muscle dense body | 93 | 21 | CC | 1.77E-05 | 1.48E-03 |
| GO:0045087 | innate immune response | 300 | 47 | BP | 5.16E-05 | 3.06E-03 |
| GO:0010332 | response to gamma radiation | 10 | 5 | BP | 7.77E-05 | 3.59E-03 |
| GO:0006869 | lipid transport | 26 | 8 | BP | 2.74E-04 | 1.12E-02 |
| GO:0098869 | cellular oxidant detoxification | 52 | 12 | BP | 5.12E-04 | 1.29E-02 |
| GO:0030017 | sarcomere | 22 | 7 | CC | 4.10E-04 | 1.29E-02 |
| GO:0006936 | muscle contraction | 17 | 6 | BP | 3.93E-04 | 1.29E-02 |
| GO:0016810 | hydrolase activity, acting on carbon-nitrogen (but not peptide) bonds | 13 | 5 | MF | 5.02E-04 | 1.29E-02 |
| GO:0071949 | FAD binding | 18 | 6 | MF | 5.94E-04 | 1.37E-02 |
| GO:0008652 | cellular amino acid biosynthetic process | 20 | 6 | BP | 1.23E-03 | 2.57E-02 |
| GO:0043050 | pharyngeal pumping | 16 | 5 | BP | 1.85E-03 | 2.96E-02 |
| GO:0016787 | hydrolase activity | 818 | 96 | MF | 2.24E-03 | 3.45E-02 |
| GO:0016311 | dephosphorylation | 63 | 12 | BP | 3.36E-03 | 3.49E-02 |
| GO:0030170 | pyridoxal phosphate binding | 48 | 10 | MF | 2.88E-03 | 3.49E-02 |
| GO:0031430 | M band | 28 | 7 | CC | 2.46E-03 | 3.49E-02 |
| GO:0003993 | acid phosphatase activity | 28 | 7 | MF | 2.46E-03 | 3.49E-02 |
| GO:0099132 | ATP hydrolysis coupled cation transmembrane transport | 23 | 6 | BP | 3.08E-03 | 3.49E-02 |

CC: cellular component; MF: molecular function; BP: biological process.

As expected there were also changes in various pathways associated with cellular stress responses (Table 1), and protein homeostasis (Table S2), with unfolded protein binding also enriched by the effect of interaction (Table S2; adj.*p* = 5.32E-05).

Taking the effect of both PD and aging into consideration, genes associated with metabolic processes, transporters of ions and lipids, and kinase activities for ATP were enriched (Table S2).

### Phenotypic effects of PD vary among genetic backgrounds

#### PD differentially affects development depending on the genetic background

Gene expression analysis indicated that PD lines were developmentally delayed. To test this directly, we scored development time, *i.e.* the time to the first appearance of eggs. Analysis of these data indicated that development was affected by PD (*p* < 2e-8), genetic background (GB; *p* < 2e-8) and the interaction between PD and GB (*p* < 0.0002). In most α-synuclein expressing lines development was delayed compared to the corresponding wild isolate (Figure 2A). NL5901 showed a very slight, yet non-significant, developmental delay compared to N2 (Figure 2A). In comparison to wild type controls, SCH1941 had the longest larval development time period and slowest reproductive development of the five transgenic lines scored (7.9 hours slower than JU1941, *p* < 1e-6, Figure 2A). SCH1926 showed an intermediate development delay of 4.3 hours slower than JU1926 (p < 0.008). SCH1511 and SCH1931 showed delay, yet not significantly (~1.8 hours and ~ 1.3 hours slower, respectively, Figure 2A) compared to the corresponding wild isolates.

**Figure 2:**
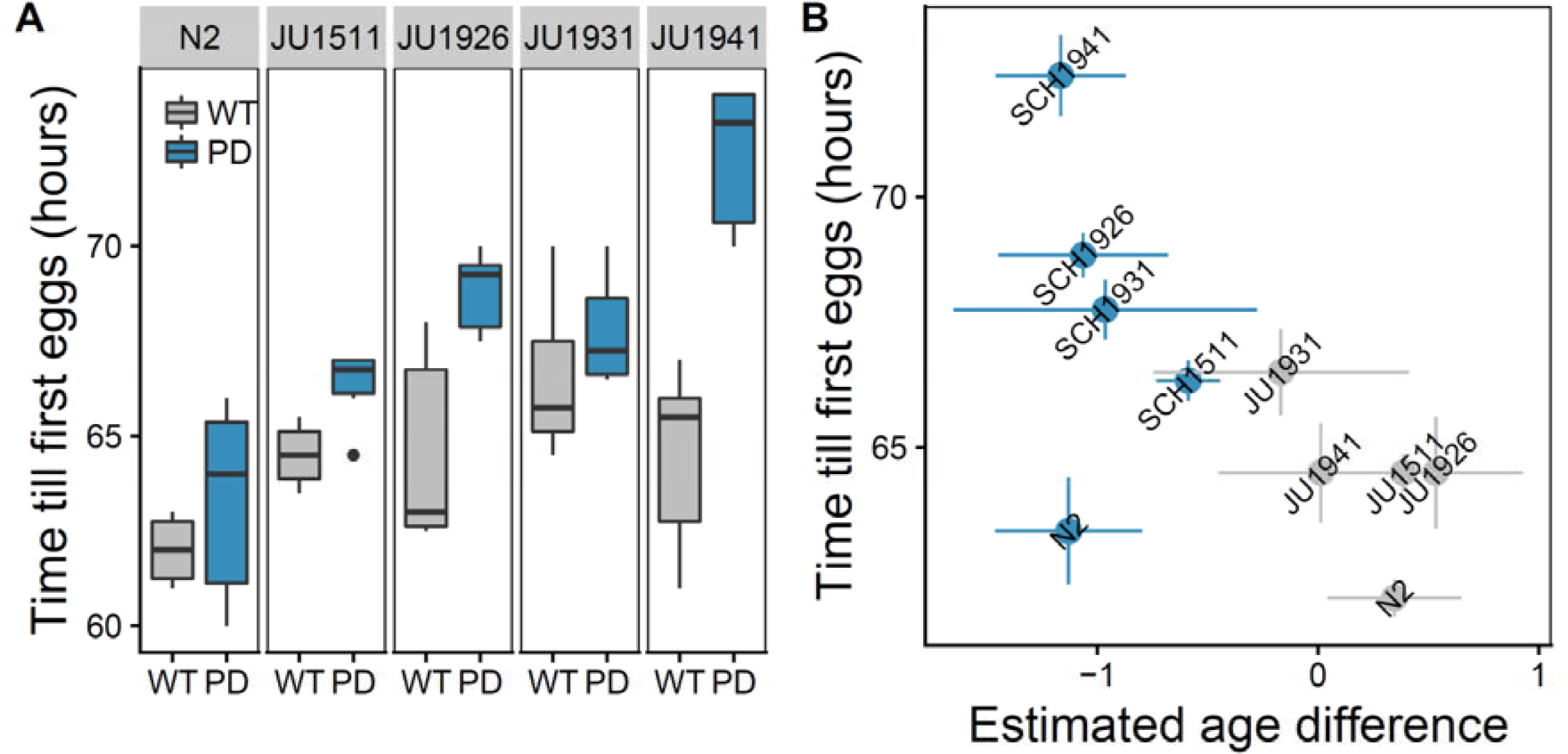
**A**) Variation in development as measured by time until first egg laying. Variation was found between the genotypes and PD introgression lines. (ANOVA: PD effect *p* < 2e-8; Genotype *p* < 2e-8; PD × Genotype Interaction *p* < 0.0002; Tuckey HSD: N2 *p* = 0.96; JU1511 *p* = 0.88; JU1926 *p* < 0.008; JU1931 *p* = 0.98; JU1941 *p* < 1e-6). **B**) Relation between estimated age at transcriptomics sampling and development time.

As found in the phenotypic assay, the transcriptomic samples of the PD lines were estimated to be younger than the corresponding wild isolate and so showed delayed development (Figure 2B, ANOVA model Estimated_Age~PD*Genotype; PD *p* < 6e-5), which was independent of the genetic background (Genotype, *p* = 0.72) and of the interaction between PD and genetic background (PD × Genotype, *p* = 0.84). Even though the ANOVA model did not show a significant effect of the genetic background and interaction on the estimated age, comparing the WT and PD lines for each individual line showed that PD1931 and JU1931 again displayed the smallest developmental difference as found in the phenotypic assay. Overall, we therefore conclude that incorporation of an α-synuclein introgression had a genotype-specific impact, differentially decreasing the developmental rate in individual genetic backgrounds.

#### PD differentially affects pumping rate depending on the genetic background

Given the observation of differential expression of genes with a function in pharyngeal pumping (Table 1), we measured the pumping rate in all lines 48 and 72 hours after recovery after L1 arrest (Figure 3). Analysis of these data found that, 72 hours after recovery from L1 arrest, pumping is affected by PD (PD, *p* < 3e-16), by the genetic background (Genotype, *p* < 3e-10) and by the interaction of PD with genetic background (PD × Genotype, *p* < 0.004; Figure 3B). Hence, the pumping rate at 72 hours in the PD lines had slowed down compared to their corresponding wild isolate (Figure 3B; N2 *p* = 0.93; JU1511 *p* < 5e-4; JU1926 *p* < 2e-5; JU1931 *p* = 0.54; JU1941 *p* < 1e-8 and comparison to Figure 3A). These data therefore indicated that the presence of an α-synuclein introgression had a noticeable genotype-specific impact on pumping rate, with these effects becoming more pronounced as worms age.

**Figure 3:**
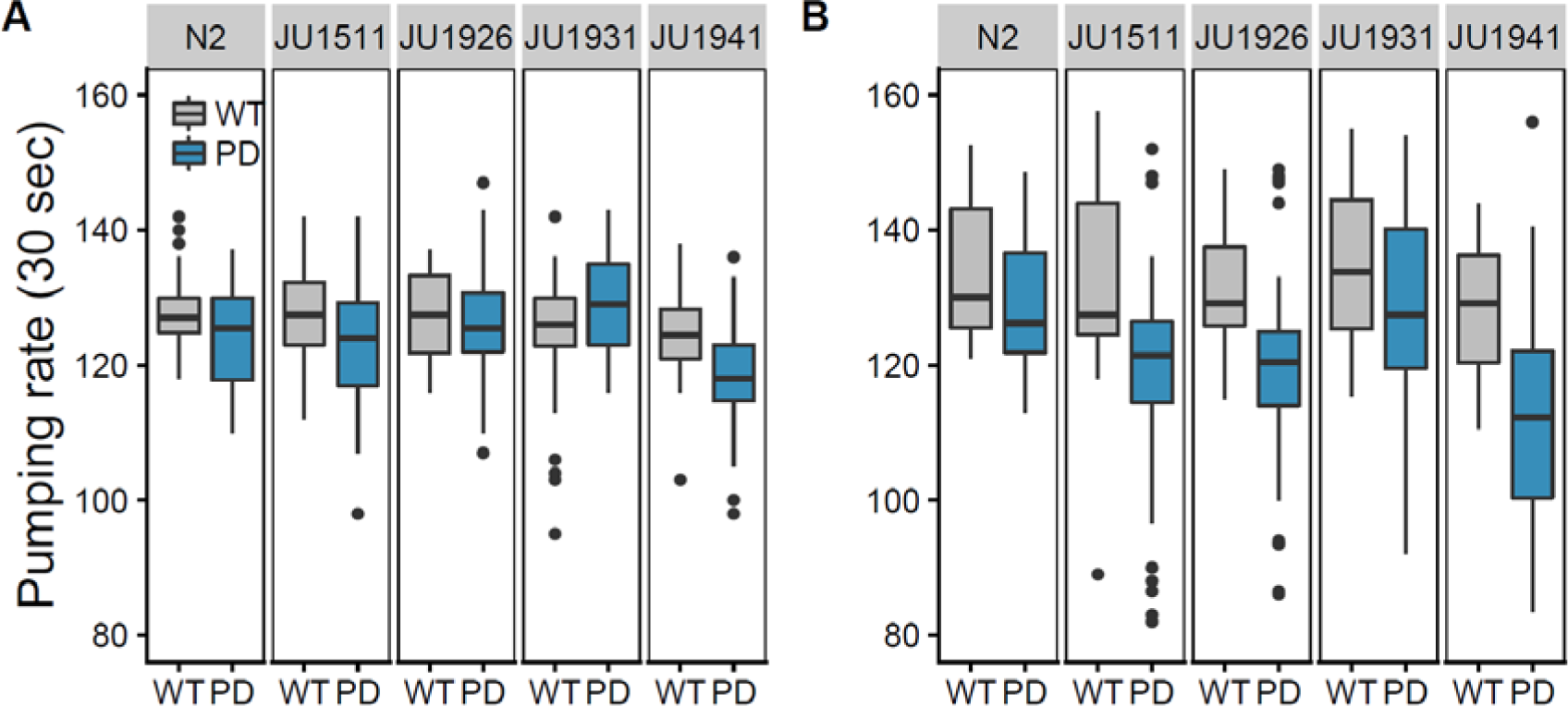
Pumping dynamics in response to PD and genetic background. Average pumping rate of 30 worms per line (A) 48 and (B) 72 hours after recovery after L1 arrest are recorded for 30 seconds. (72 hour old worms ANOVA: PD *p* < 1e-16; Genetic background *p* < 3e-10; Interaction *p* < 0.0004; Tucky HSD: N2 *p* = 0.93; JU1511 *p* < 5e-4; JU1926 *p* < 2e-5; JU1931 *p* = 0.54; JU1941 *p* < 1e-8).

#### PD decreases lifespan in some genetic backgrounds

Given the age-related changes in pumping rate and enrichment of genes associated with aging and stress response pathways in response to PD, we hypothesised that the introgressed PD transgene could affect longevity. Therefore, we measured lifespan in all lines. Comparison of N2 and NL5901 (PD in an N2 background) showed that lifespan was not affected by PD in the N2 genetic background (Figure 4; N2 log-rank *p* = 0.14, mean age *p* = 0.99). However, in the wild isolate genetic backgrounds, lines containing the **α**-synuclein introgression displayed significantly accelerated death (Figure 4A) and a shortened lifespan (Figure 4B). These data also indicated that the **α**-synuclein introgression in the wild isolate genetic backgrounds resulted in increased rates of maternal hatching (bagging) (Table S3).

**Figure 4:**
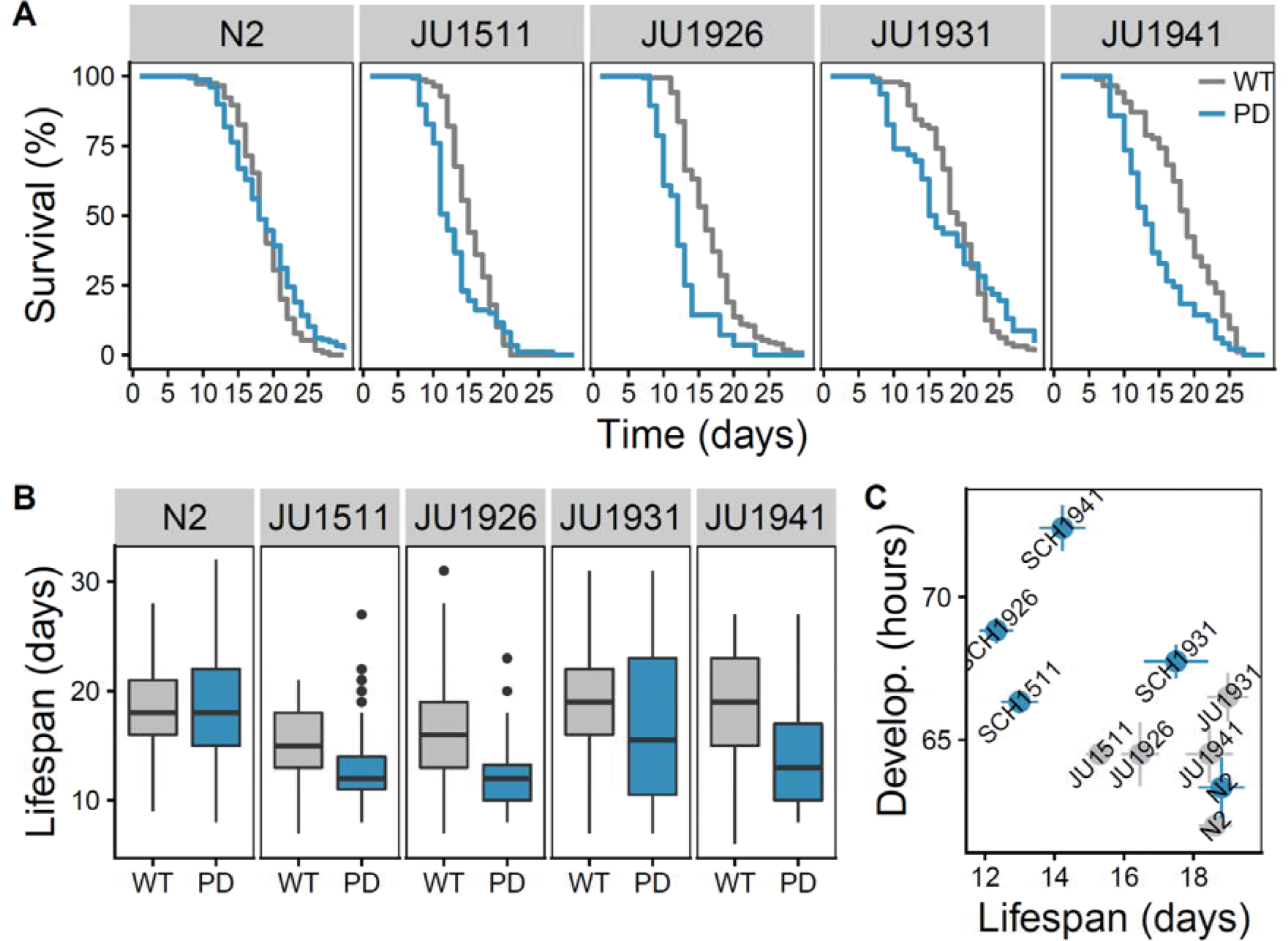
*C. elegans* PD ILs constitutively expressing full-length human α-synuclein exhibit significantly shorter lifespan. (**A**) Survival curves for wild-type (WT) and transgenic nematodes expressing α-synuclein (PD) (log-rank test (‘survival’ package, R): N2 *p* = 0.14; JU1511 *p* < 0.003; JU1926 *p* < 4*1e-7; JU1931 *p* = 0.79 (first 20 days *p* < 0.0004); JU1941 *p* < 7e-5). (**B**) Mean lifespan for WT and PD lines (ANOVA: PD effect *p* < 3e-5; Genetic background *p* << 1e-16; Interaction *p* < 0.0002; Tucky HSD: N2 *p* = 0.99; JU1511 *p* < 0.008; JU1926 *p* < 0.0004; JU1931 *p* = 0.71; JU1941 *p* < 1e-5). **C**) Relation between lifespan at life history sampling and development time.

This may represent a failure of the vulval muscles associated with egg-laying as a result of α-synuclein expression and aggregation. These observations indicate that α-synuclein expression lowers lifespan and organismal fitness in some genotypes, but not others.

#### Genetic background effects PD outcome

Comparison of phenotypic effects across the genetic backgrounds tested, suggested that the phenotypes associated with the α-synuclein transgene introgression vary between genotypes in a consistent manner (Figure 5). Multiple phenotypes were affected by PD in the JU1511, JU1926, and JU1941 background, whereas they were less affected in the JU1931 and N2 backgrounds. This suggests a shared genetic component across many of the phenotypes we have observed, but does also suggest some genotype-specific variation, *e.g.* no pair of lines shows the same pattern of phenotypic effects (Figure 5).

**Figure 5:**
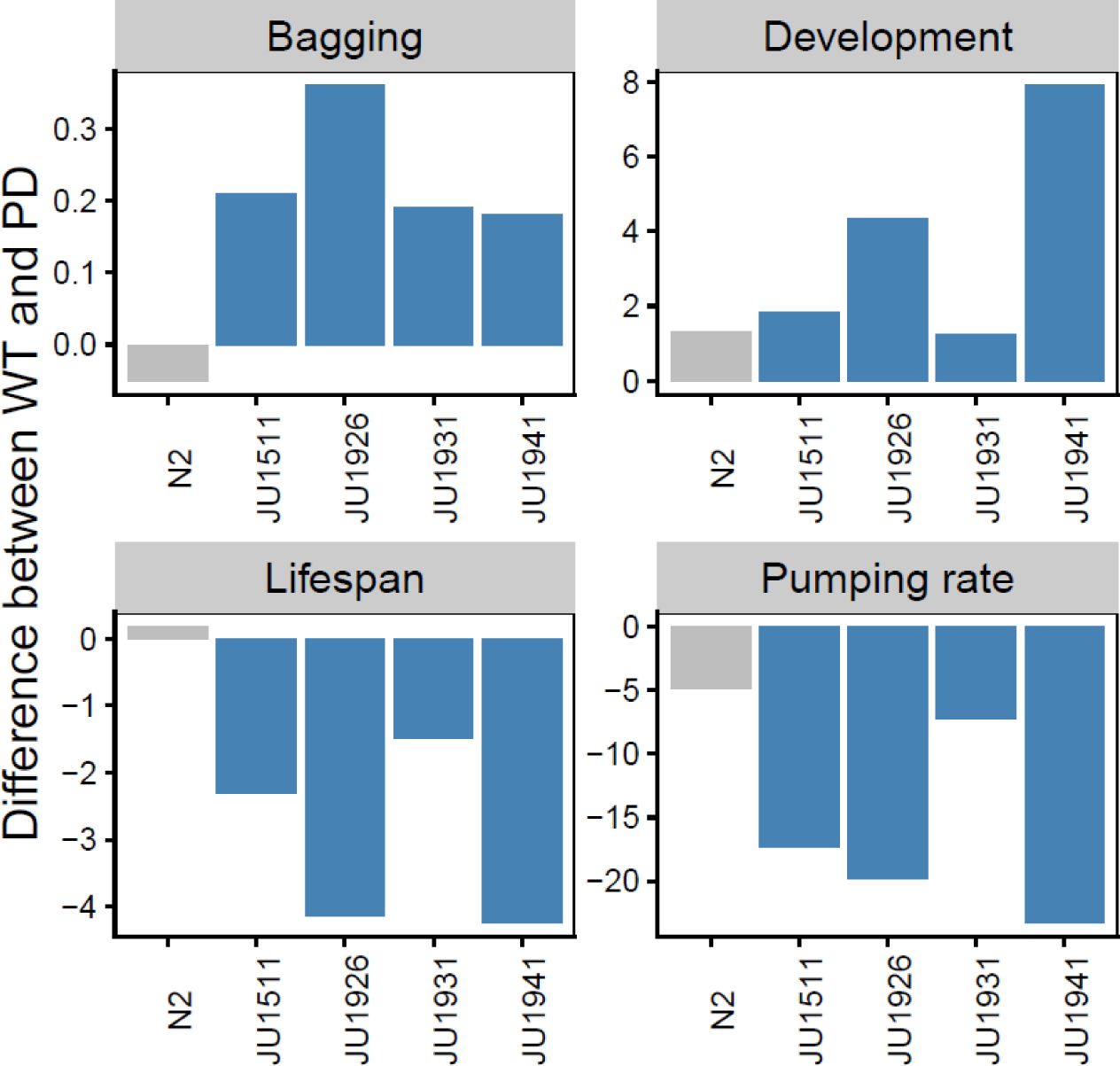
Phenotypic effects of α-synuclein expression across different genetic backgrounds. Differences between wild type and α-synuclein expressing lines in bagging (% difference in matricidal hatching), in development (difference in hours from L1 arrest to first egg lay), in lifespan (difference in mean lifespan in days) and in pumping rate (difference in number of pumps recorded in a 30s period).

As alleles from different *C. elegans* genomes can produce a range of synthetic deleterious effects resulting in full or partially genomic incompatibilities between their genomes (Seidel *et al.* 2008; Snoek *et al.* 2014b), we sought to test if the N2 region alone replicated the phenotypes we observed here. We therefore introgressed the *pkIs2386* transgene into a CB4856 genetic background, generating the SCH4856 line and undertook comparisons of this line with CB4856 and CBN93, a line with an introgression of the N2 genome spanning the 3.3-12.8Mbp region on chromosome IV in an CB4856 background (Table S1). This control was undertaken as deleterious interactions between alleles from N2 and those from CB4856 are well characterized (Seidel *et al.* 2008; Snoek *et al.* 2014b). Comparisons of pumping rate and of development time between these lines indicated that the expression of α-synuclein in SCH4856 produced effects not seen in CBN93 (Figure 6) and that these effects mirror those seen in the other genetic backgrounds. This provides strong support for the view that the phenotypic effects we observe were a consequence of α-synuclein expression and not the introgression of the N2 region.

**Figure 6:**
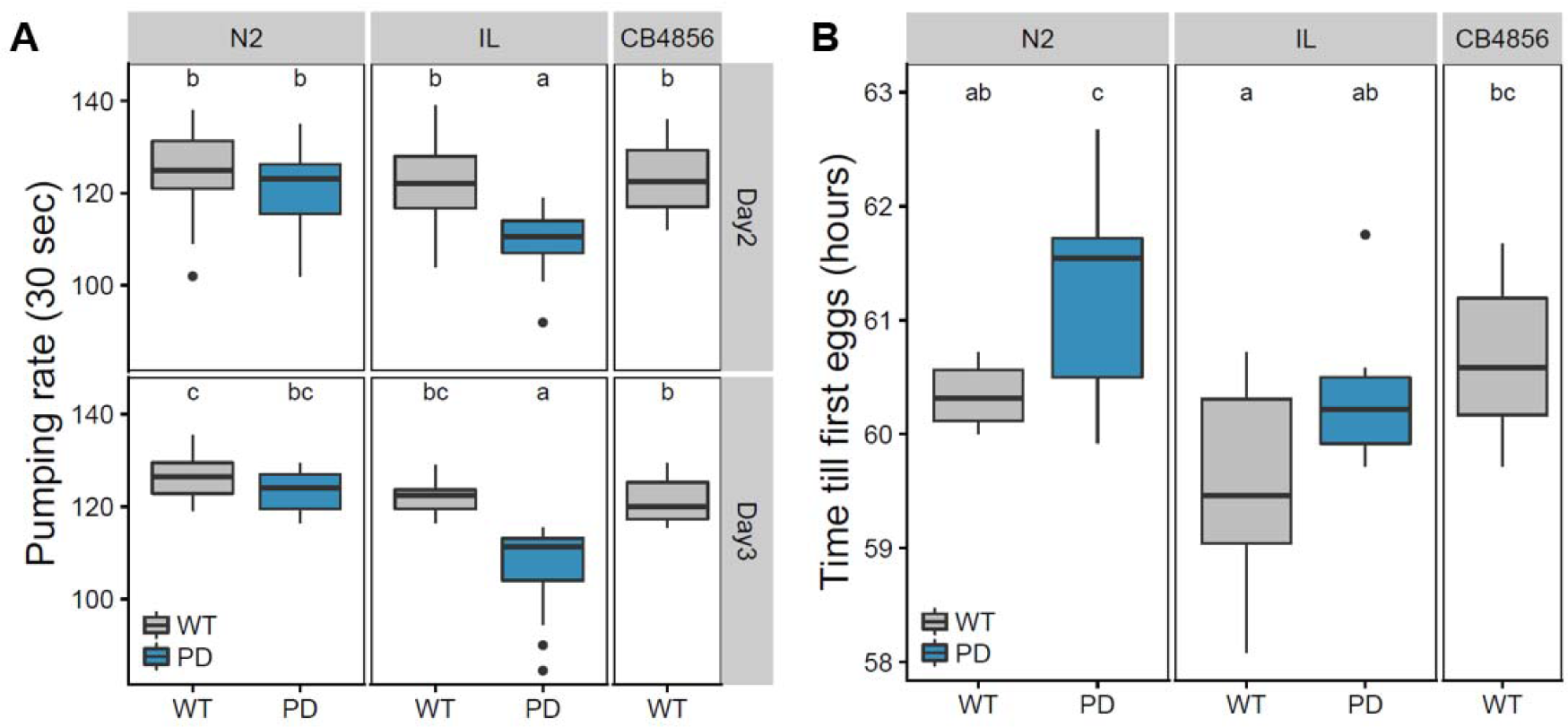
Phenotypic effects observed in a CB4856 genetic background depend on α-synuclein expression rather than N2 alleles. (**A**) Average pumping rate in 30s of worms 48 and 72 hours after recovery after L1 arrest, and (**B**) Time from recovery of L1 arrest to first egg lay. Letter codes denote lines that differ significantly (*p* < 0.05).

## Discussion

We have introgressed a transgene that results in the expression of α-synuclein in the body wall muscle into different genetic backgrounds of *C. elegans.* Analysis of these newly created α-synuclein ILs indicates that genetic background effects both the response of the animals to α-synuclein expression at the level of gene expression and in terms of the phenotypic consequences. Our gene expression analysis identifies a range of changes associated with α-synuclein expression. Many of these - particularly those involved in muscle function and in various stress responses - matched expectations given the site of expression and the known effects of α-synuclein aggregation on cellular function. However we also identify changes in lipid metabolism (Table 1), an important result given that this has been previously linked to α-synuclein pathology in a yeast model (Cooper *et al.* 2006) and that there is evidence of a direct association between α-synuclein and lipid droplets (Cole *et al.* 2002). At a phenotypic level, changes in lipid levels in a *C. elegans* model of PD have also been observed (Jadiya *et al.* 2011), our results therefore provide a set of candidate genes to investigate how this is mediated and also show that such effects can be expected to vary across genetic backgrounds.

On a phenotypic level we found that, in comparison to corresponding wild isolates, the PD lines: developed more slowly; suffered an increased rate of decline in pharyngeal pumping; exhibited a shorter lifespan; and died from matricidal hatching of eggs at an increased rate. Similar phenotypic effects are observed in other *C. elegans* models of protein misfolding disease. For example, transgenic worms expressing polyQ proteins and Aβ both show a significantly shortened lifespan (Gidalevitz *et al.* 2013; Fong *et al.* 2016). A reduced lifespan could therefore be a general toxicity phenotype, providing an indirect measure of organismal dysfunction caused by misfolded protein aggregations in the body wall. In combination with the observed changes in various metabolic processes, this may also indicate crosstalk between PD transgene and genetic backgrounds. For example, the autophagy process plays a role in lipid storage in *C. elegans*, which is also determined to be key in mitochondrial dysfunctions in neurodegenerative disorders (Lapierre *et al.* 2012; Singh *et al.* 2016).

Many of the phenotypic changes we see are either only found in wild isolate genetic backgrounds or are more pronounced in these backgrounds (Table 2). This demonstrates that there is variation between isolates that appears to affect different stages of the aggregation process. Previously, only a single *C. elegans* study has directly investigated the effect of genetic background in the context of protein misfolding disease. That study found complex variation in polyglutamine (polyQ40) aggregation and toxicity in three wild isolate backgrounds and in a panel of 21 recombinant inbred lines (RILs) (Gidalevitz *et al.* 2013). The RIL analysis also showed that various pathological effects could be uncoupled (Gidalevitz *et al.* 2013). In this study, we also show some evidence that natural genetic variation in *C. elegans* can uncouple different effects of α-synuclein aggregation (Table 2), suggesting that this may be a general pattern.

A caveat with this interpretation is however that a range of incompatibilities between alleles from different *C. elegans* genomes have been identified (Seidel *et al.* 2008; Snoek *et al.* 2014b) and that the region surrounding the site of the α-synuclein transgene contains a number of mapped quantitative trait loci affecting various life history traits. For example, ILs with introgressed regions of chromosome IV from CB4856 in an N2 genetic background have identified QTLs affecting lifespan and pumping rate that partially overlap our introgression (Doroszuk *et al.* 2009). Similarly, a complex interaction between alleles from the N2 and CB4856 genomes has been found on chromosome IV (Snoek *et al.* 2014b). However, little is known about the frequency of such synthetic deleterious effects between other genotypes of *C. elegans.* Proteomic analysis of age-dependent changes in protein solubility by Rodriguez *et al.* (2012) also indicates that the genes encoding proteins that become insoluble with age are enriched for modifiers of lifespan. Additionally, a heat-stress specific QTL for recovery (Rodriguez *et al.* 2012) as well as an eQTL-hotspot have been identified on the left arm of chromosome IV (IV: 1.0-2.5 Mb) (Snoek *et al.* 2017), while a QTL for maternal hatching rate is at the position of the introgression (chromosome IV ~6M) (Stastna *et al.* 2015). That such effects might alter disease-related processes complicates the interpretation of both our results here and the previous work of Gidalevitz *et al.* (2013). To experimentally address this caveat, we introgressed the α-synuclein transgene into a CB4856 genetic background and undertook comparisons with a line that contains a comparable region of the N2 genome, but no transgene, in a CB4856 background. These comparisons indicated that the N2 introgression alone does not recapitulate the phenotypic effects seen in the α-synuclein expressing line (Figure 6). As these comparisons again show that the expression of α-synuclein in a CB4856 genetic background results in different and more severe effects than those seen in an N2 background, this strongly supports the view that genetic background needs to be considered in the case of PD. A move to the use of more defined modifications in a range of genetic backgrounds that limited the introduction of other alleles - perhaps via the use of CRISPR to introduce the relevant transgenes - would therefore be ideal. As there is extensive variation between isolates of *C. elegans* in lifespan and in various stress responses and in lifespan, and that variation has now been identified for two protein misfolding diseases such a development would facilitate the systematic analysis of the role of natural variation in such diseases.

## Methods

### *C. elegans* lines and growth conditions

Worms were maintained at 20°C on nematode growth medium (NGM) plates seeded with *Escherichia coli* 0P50 (Brenner 1974). To obtain starved and synchronized L1 larvae, gravid adults were bleached to isolate eggs, which then were allowed to hatch overnight at 20°C (Brenner 1974). The *C. elegans* wild types JU1511, JU1926, JU1931, JU1941, canonical strain N2 (Bristol), and transgenic strain NL5901, which contains the *pkIs2386* transgene [unc-54p::α-synuclein::YFP + unc-119(+)], were obtained from the Caenorhabditis Genetics Center. To introgress the α-synuclein::YFP transgene into the wild backgrounds, NL5901 hermaphrodites were mated with wild males, then five to ten F1 fluorescent males were mated with wild hermaphrodites for F2 fluorescent males. This backcrossing to the wild background was repeated until generation F7 was reached. At this point, strains were allowed to self-fertilise to obtain homozygotes lines and then cryopreserved.

### IL genotyping

#### PCR-based Genotyping

According to the different expressions of DNA hybridizations induced marker genes between N2 and the four wild isolates (Figure S1), we performed PCR-based genotyping on chromosome IV with N2, wild isolates and all five PD lines. DNA was isolated from the lysates of 5 individual adults each line, and then was used for genotyping PCRs. Genotyping primers utilized insertions/deletions between the CB4856 and N2 genomes, with 41 primer pairs covering the genome used (see Table S1 for details of primers and marker locations). PCR was carried out with the GoTaq DNA polymerase kit (Promega) according to the manufacturer’s recommendations. The sizes of amplified products were assessed by electrophoresis in 1.5% agarose gels stained with Ethidium Bromide.

#### Hybridization markers

Marker genes found by DNA hybridizations in Volkers *et al.* (2013) were used to find the genomic position of the PD introgression in each genetic background. We tested which genomic region was missing markers for the wildtype background, this region then must be from the NL5901 background.

### Sample preparations and RNA microarray analysis

#### mRNA microarrays

Three independent replicates of each strain (synchronized 48hrs old L4 larvae) were analyzed. Briefly, worms were generated by hatching alkaline hypochlorite-purified eggs and then harvested by centrifugation, washed with M9 buffer, frozen in liquid N2, and stored at -80 °C until use. The Maxwell^®^ 16 Tissue LEV Total RNA Purification Kit was used for mRNA isolation, following the manufacturer’s protocol with a modified lysis step. In addition to the lysis buffer, proteinase K was added and the sample was incubated at 65^°^C while shaking at 1000 rpm for 10 minutes. Thereafter the standard protocol was followed. PolyA RNA was used to generated Cy3 and Cy5-labeled cRNA samples, which were then hybridized to 4X44K slides V2 (Agilent) *C. elegans* whole genome GeneChips, processed, and scanned (full microarray data in ArrayExpress with accession E-MTAB-6960). RNA microarray statistical analysis and data processing were performed using the Limma package for the R software environment (https://www.bioconductor.org/packages/release/bioc/html/limma.html). To find the genes affected by PD, genotype and age, these terms were used as explanatory factors in a linear model (gene expression ~ age + PD * genotype). Significance thresholds were determined by permutations of all spots on the array. In the permutations, the RNA hybridization intensities were randomly distributed over the genotypes and batches. Therefore, the p-value that gave a min ratio of false positives/true positives of 0.05 (= -log10(p) > 3.4) was set for convenience, *i.e.* an FDR at 0.0186 for PD effect, 0.0249 for genotype effect and 0.073 for the interaction between PD and genotype.

#### Enrichment

Enrichments were done on genes groups divined by their significance from the linear model a -log10(p) > 2 was used, then a hyper geometric test in R was used to test each GO term for enrichment.

#### PCA

To partition the variation in gene expression a PCA was done on the transcription profiles (the log2 ratios with the mean) of all samples. The first two axis were used for visualisation.

#### Age estimation

Age estimation of the worms sampled for transcriptomics was done by comparing the class I age responsive genes from Snoek *et al.* (2014a), as done by Snoek *et al.* 2014a; van der Bent *et al.* 2014; Jovic *et al.* 2017.

### Phenotypic assays

#### Lifespan

Nematodes were cultured on 0P50 bacteria without fluorouracil deoxyribose (FUdR), and transferred away from their progeny every day during the vigorous reproductive period and every other day during the reduced reproductive period. From the young adult stage, worms were examined for signs of life daily. Individuals that were not moving or twitching after gentle stimulation, followed by vigorous stimulation, or that did not exhibit pharyngeal pumping for 30 seconds, were considered dead. Individual worms that had died from internal hatching of progeny, or that had crawled off the plates, were censored from lifespan result but were used for maternal hatching analysis. Data analysis was performed in R using the ‘survival’ package.

#### Development time

Synchronised L1 juveniles were obtained by allowing eggs to hatch in M9 buffer after three washes in M9 after hypochlorite treatment. Arrested L1s were then transferred to NGM plates, which fed with *E. coli* 0P50 and were incubated at 20°C. Tracking observations and inspections were done at regular time intervals. Development time was defined as the period between worm inoculation and the moment at which the first appearance of eggs and the period until the reproductions reach peak level.

#### Pharyngeal pumping

Synchronized individual worms were observed 48 hours and 72 hours after bleaching. The number of contractions in the terminal bulb of pharynx was counted for 30s (n = 19) for the 48 hour-old worms, then counting increased to 1 minute for 3 day-old worms (n = 30).

#### Statistical analysis

Phenotypic differences between the lines were tested using an ANOVA (model: phenotype~PD*GB) to test for overall effects and Tukey HSD to test for lines specific differences. Survival curves were tested by log-rank test from the “survival” package in R.

## Declarations

Ethics approval and consent to participate - Not applicable

Consent for publication - Not applicable

## Availability of data and material

All data generated or analysed during this study are included in this published article [and its supplementary information files].

## Competing interests

The authors declare that they have no competing interests.

## Funding

This work is supported by a Leverhulme Trust Research Grant awarded to SH, and by BLS NWO funding awarded to JK. Some strains were provided by the CGC, which is funded by NIH Office of Research Infrastructure Programs (P40 0D010440). These funding bodies had no role in the design of the study, the collection, analysis, and interpretation of data, or in writing the manuscript.

## Authors’ contributions

YW constructed the introgression lines and undertook the phenotypic analyses with assistance from JS. YW and JR performed the microarray. YW, LBS and MGS analysed the data. SH, JK and YW conceived the project. SH, YW, LBS and JK wrote the manuscript. All authors read and approved the final manuscript.

## Acknowledgements

We thank Yu Nie for comments and assistance with assays. We also thank the CGC for worm strains.

## Supplementary material

Additional file 1: Microarray data (“Pvalues_PD_paper.txt”)

Table S1 : Markers used in the genotyping of PD ILs.

Table S2: Gene ontology analysis (GO) of genes identified as affected by Age, Genotype, PD and the interaction between genotype and PD.

Table S3: Rates of matricidal hatching in PD ILs and relevant wild isolates.

Figure S1: Gene expression markers across the genome. Determination of the PD introgression in the four wild isolate backgrounds. Marker genes with different expression between N2 and the wild isolates (All) were used to detect the PD introgression and N2 border regions. The marker genes missing in the PD lines indicate the N2 border regions and position of the PD introgression (PD Missing). Different genetic backgrounds are indicated by the different colours. The position(s) where all lines have missing markers show the likely PD locus, the extra missing markers on chromosome V show a possible extra introgression in SCH1931.

## References

Aarsland D, Kurz MW. The epidemiology of dementia associated with Parkinson disease. Journal of the Neurological Sciences. 2010;289(1):18–22.

Aybek S, Lazeyras F, Gronchi-Perrin A, Burkhard PR, Villemure JG, Vingerhoets FJ. Hippocampal atrophy predicts conversion to dementia after STN-DBS in Parkinson’s disease. Parkinsonism & Related Disorders. 2009;15(7):521–4.

van der Bent ML, Sterken MG, Volkers RJ, Riksen JA, Schmid T, Hajnal A, Kammenga JE, Snoek LB. Loss-of-function of β-catenin bar-1 slows development and activates the Wnt pathway in *Caenorhabditis elegans*. Scientific Reports. 2014;4:4926.

Brenner S. The genetics of *Caenorhabditis elegans*. Genetics. 1974;77(1):71–94.

Calahorro F, Ruiz-Rubio M. *Caenorhabditis elegans* as an experimental tool for the study of complex neurological diseases: Parkinson’s disease, Alzheimer’s disease and autism spectrum disorder. Invertebrate Neuroscience. 2011;11(2):73–83.

Coe TS, Hamilton PB, Griffiths AM, Hodgson DJ, Wahab MA, Tyler CR. Genetic variation in strains of zebrafish *(Danio rerio)* and the implications for ecotoxicology studies. Ecotoxicology. 2009;18(1):144–50.

Cole NB, Murphy DD, Grider T, Rueter S, Brasaemle D, Nussbaum RL. Lipid droplet binding and oligomerization properties of the Parkinson’s disease protein α-synuclein. Journal of Biological Chemistry. 2002;277(8):6344–52.

Cooper AA, Gitler AD, Cashikar A, Haynes CM, Hill KJ, Bhullar B, Liu K, Xu K, Strathearn KE, Liu F, Cao S. **α**-Synuclein blocks ER-Golgi traffic and Rab1 rescues neuron loss in Parkinson’s models. Science. 2006;313(5785):324–8.

Doroszuk A, Snoek LB, Fradin E, Riksen J, Kammenga J. A genome-wide library of CB4856/N2 introgression lines of *Caenorhabditis elegans*. Nucleic Acids Research. 2009;37(16):e110–.

Duveau F, Félix MA. Role of pleiotropy in the evolution of a cryptic developmental variation in *Caenorhabditis elegans*. PLoS Biology. 2012;10(1):e1001230.

Elvin M, Snoek LB, Frejno M, Klemstein U, Kammenga JE, Poulin GB. A fitness assay for comparing RNAi effects across multiple *C. elegans* genotypes. BMC Genomics. 2011;12(1):510.

Fong S, Teo E, Ng LF, Chen CB, Lakshmanan LN, Tsoi SY, Moore PK, Inoue T, Halliwell B, Gruber J. Energy crisis precedes global metabolic failure in a novel *Caenorhabditis elegans* Alzheimer disease model. Scientific Reports. 2016;6:33781.

Fonte V, Kipp DR, Yerg J, Merin D, Forrestal M, Wagner E, Roberts CM, Link CD. Suppression of in vivo β-amyloid peptide toxicity by overexpression of the HSP-16.2 small chaperone protein. Journal of Biological Chemistry. 2008;283(2):784–91.

Gaertner BE, Phillips PC. *Caenorhabditis elegans* as a platform for molecular quantitative genetics and the systems biology of natural variation. Genetics research. 2010;92(5-6):331–48.

Gidalevitz T, Wang N, Deravaj T, Alexander-Floyd J, Morimoto RI. Natural genetic variation determines susceptibility to aggregation or toxicity in a *C. elegans* model for polyglutamine disease. BMC Biology. 2013;11(1):100.

Gratwicke J, Jahanshahi M, Foltynie T. Parkinson’s disease dementia: a neural networks perspective. Brain. 2015;138(6):1454–76.

van Ham TJ, Thijssen KL, Breitling R, Hofstra RM, Plasterk RH, Nollen EA. *C. elegans* model identifies genetic modifiers of α-synuclein inclusion formation during aging. PLoS Genetics. 2008;4(3):e1000027.

Jadiya P, Khan A, Sammi SR, Kaur S, Mir SS, Nazir A. Anti-Parkinsonian effects of *Bacopa monnieri:* insights from transgenic and pharmacological *Caenorhabditis elegans* models of Parkinson’s disease. Biochemical and Biophysical Research Communications. 2011;413(4):605–10.

Jovic K, Sterken MG, Grilli J, Bevers RP, Rodriguez M, Riksen JA, Allesina S, Kammenga JE, Snoek LB. Temporal dynamics of gene expression in heat-stressed *Caenorhabditis elegans*. PloS One. 2017;12(12):e0189445.

Justice MJ, Dhillon P. Using the mouse to model human disease: increasing validity and reproducibility. Disease Models & Mechanisms 2016;9:101–103.

Kamkina P, Snoek LB, Grossmann J, Volkers RJ, Sterken MG, Daube M, Roschitzki B, Fortes C, Schlapbach R, Roth A, von Mering C. Natural genetic variation differentially affects the proteome and transcriptome in *Caenorhabditis elegans*. Molecular & Cellular Proteomics. 2016;15(5):1670–80.

Kammenga JE, Phillips PC, De Bono M, Doroszuk A. Beyond induced mutants: using worms to study natural variation in genetic pathways. Trends in Genetics. 2008;24(4):178–85.

Kammenga JE. The background puzzle: how identical mutations in the same gene lead to different disease symptoms. The FEBS journal. 2017; 284:3362–3373.

Kearney JA. Genetic modifiers of neurological disease. Current Opinion in Genetics & Development. 2011;21(3):349–53.

Kivipelto M, Ngandu T, Fratiglioni L, Viitanen M, Kåreholt I, Winblad B, Helkala EL, Tuomilehto J, Soininen H, Nissinen A. Obesity and vascular risk factors at midlife and the risk of dementia and Alzheimer disease. Archives of Neurology. 2005;62(10):1556–60.

Lapierre LR, Meléndez A, Hansen M. Autophagy links lipid metabolism to longevity in *C. elegans*. Autophagy. 2012;8(1):144–146.

Li Y, Álvarez OA, Gutteling EW, Tijsterman M, Fu J, Riksen JA, Hazendonk E, Prins P, Plasterk RH, Jansen RC, Breitling R. Mapping determinants of gene expression plasticity by genetical genomics in *C. elegans*. PLoS Genetics. 2006;2(12):e222.

Martin N, Singh J, Aballay A. Natural genetic variation in the *Caenorhabditis elegans* response to *Pseudomonas aeruginosa*. G3: Genes, Genomes, Genetics. 2017;7(4):1137–47.

McColl G, Roberts BR, Pukala TL, Kenche VB, Roberts CM, Link CD, Ryan TM, Masters CL, Barnham KJ, Bush AI, Cherny RA. Utility of an improved model of amyloid-beta 142) toxicity in *Caenorhabditis elegans* for drug screening for Alzheimer’s disease. Molecular Neurodegeneration. 2012;7(1):57.

Reynolds RM, Phillips PC. Natural variation for lifespan and stress response in the nematode *Caenorhabditis remanei*. PLoS One. 2013;8(4):e58212.

Rockman MV, Skrovanek SS, Kruglyak L. Selection at linked sites shapes heritable phenotypic variation in *C. elegans*. Science. 2010;330(6002):372–6.

Rodriguez M, Snoek LB, Riksen JA, Bevers RP, Kammenga JE. Genetic variation for stress-response hormesis in *C. elegans* lifespan. Experimental gerontology. 2012;47(8):581–7.

Rodriguez M, Snoek LB, De Bono M, Kammenga JE. Worms under stress: *C. elegans* stress response and its relevance to complex human disease and aging. Trends in Genetics. 2013;29(6):367–74.

Schmid T, Snoek LB, Fröhli E, van der Bent ML, Kammenga J, Hajnal A. Systemic regulation of RAS/MAPK signaling by the serotonin metabolite 5-HIAA. PLoS Genetics. 2015;11(5):e1005236.

Schulz-Schaeffer WJ. The synaptic pathology of α-synuclein aggregation in dementia with Lewy bodies, Parkinson’s disease and Parkinson’s disease dementia. Acta Neuropathologica. 2010;120(2):131–43.

Seelaar H, Kamphorst W, Rosso SM, Azmani A, Masdjedi R, de Koning I, Maat-Kievit JA, Anar B, Kaat LD, Breedveld GJ, Dooijes D. Distinct genetic forms of frontotemporal dementia. Neurology. 2008;71(16):1220–6.

Seidel HS, Rockman MV, Kruglyak L. Widespread genetic incompatibility in *C. elegans* maintained by balancing selection. Science. 2008;319(5863):589–94.

Sin O, Michels H, Nollen EA. Genetic screens in *Caenorhabditis elegans* models for neurodegenerative diseases. Biochimica et Biophysica Acta (BBA)-Molecular Basis of Disease. 2014;1842(10):1951–9.

Singh KD, Roschitzki B, Snoek LB, Grossmann J, Zheng X, Elvin M, Kamkina P, Schrimpf SP, Poulin GB, Kammenga JE, Hengartner MO. Natural genetic variation influences protein abundances in *C. elegans* developmental signalling pathways. PloS One. 2016;11(3):e0149418.

Singleton AB, Farrer M, Johnson J, Singleton A, Hague S, Kachergus J, Hulihan M, Peuralinna T, Dutra A, Nussbaum R, Lincoln S. α-Synuclein locus triplication causes Parkinson’s disease. Science. 2003;302(5646):841.

Snoek BL, Sterken MG, Bevers RP, Volkers RJ, van’t Hof A, Brenchley R, Riksen JA, Cossins A, Kammenga JE. Contribution of trans regulatory eQTL to cryptic genetic variation in *C. elegans*. BMC genomics. 2017;18(1):500.

Snoek LB, Sterken MG, Volkers RJ, Klatter M, Bosman KJ, Bevers RP, Riksen JA, Smant G, Cossins AR, Kammenga JE. A rapid and massive gene expression shift marking adolescent transition in *C. elegans*. Scientific Reports. 2014a;4:3912.

Snoek LB, Orbidans HE, Stastna JJ, Aartse A, Rodriguez M, Riksen JA, Kammenga JE, Harvey SC. Widespread genomic incompatibilities in *Caenorhabditis elegans*. G3: Genes, Genomes, Genetics. 2014b;4(10):1813–23.

Stastna JJ, Snoek LB, Kammenga JE, Harvey SC. Genotype-dependent lifespan effects in peptone deprived *Caenorhabditis elegans*. Scientific reports. 2015;5:16259.

Steinkraus KA, Smith ED, Davis C, Carr D, Pendergrass WR, Sutphin GL, Kennedy BK, Kaeberlein M. Dietary restriction suppresses proteotoxicity and enhances longevity by an *hsf-1*-dependent mechanism in *Caenorhabditis elegans*. Aging Cell. 2008;7(3):394–404.

Sterken MG, van der Plaat LV, Riksen JA, Rodriguez M, Schmid T, Hajnal A, Kammenga JE, Snoek BL. Ras/MAPK modifier loci revealed by eQTL in *Caenorhabditis elegans*. G3: Genes, Genomes, Genetics. 2017;7(9):3185–93.

Teschendorf D, Link CD. What have worm models told us about the mechanisms of neuronal dysfunction in human neurodegenerative diseases? Molecular Neurodegeneration. 2009;4(1):38.

Viñuela A, Snoek LB, Riksen JA, Kammenga JE. Genome-wide gene expression regulation as a function of genotype and age in *C. elegans*. Genome Research. 2010;20(7):929–37.

Volkers RJ, Snoek LB, van Hellenberg Hubar CJ, Coopman R, Chen W, Yang W, Sterken MG, Schulenburg H, Braeckman BP, Kammenga JE. Gene-environment and protein-degradation signatures characterize genomic and phenotypic diversity in wild *Caenorhabditis elegans* populations. BMC Biology. 2013;11(1):93.

Wade CM, Daly MJ. Genetic variation in laboratory mice. Nature Genetics. 2005;37(11):1175.

Wang YA, Kammenga JE, Harvey SC. Genetic variation in neurodegenerative diseases and its accessibility in the model organism *Caenorhabditis elegans*. Human Genomics. 2017;11(1):12.

